# Temporal patterns of vampire bat rabies and host connectivity in Belize

**DOI:** 10.1101/2020.07.16.204446

**Authors:** Daniel J. Becker, Alice Broos, Laura M. Bergner, Diana K. Meza, Nancy B. Simmons, M. Brock Fenton, Sonia Altizer, Daniel G. Streicker

## Abstract

In the Neotropics, vampire bats (*Desmodus rotundus*) are the main reservoir host for rabies, a highly fatal encephalitis caused by viruses in the genus *Lyssavirus*. Although patterns of rabies virus exposure and infection have been well-studied for vampire bats in South America and Mexico, exploring the ecology of vampire bat rabies in other regions is crucial for predicting risks to livestock and humans. In Belize, rabies outbreaks in livestock have increased in recent years, underscoring the need for systematic data on viral dynamics in vampire bats. In this study, we examine the first three years of a longitudinal study on the ecology of vampire bat rabies in northern Belize. Rabies seroprevalence in bats was high across years (29–80%), suggesting active and endemic virus circulation. Across two locations, the seroprevalence time series per site were inversely related and out of phase by at least a year. Microsatellite data demonstrated historic panmixia of vampire bats, and mark–recapture detected rare but contemporary inter-site dispersal. This degree of movement could facilitate spatial spread of rabies virus but is likely insufficient to synchronize infection dynamics, which offers one explanation for the observed phase lag in seroprevalence. More broadly, our analyses suggest frequent transmission of rabies virus within and among vampire bat roosts in northern Belize and highlight the need for future spatiotemporal, phylogenetic, and ecological studies of vampire bat rabies in Central America.

## Introduction

Rabies is an acute, highly lethal encephalitis caused by viruses in the family *Rhabdoviridae*, genus *Lyssavirus* (Wunner and Jackson, 2010). Lyssaviruses most likely originated from bats (Order Chiroptera) (Badrane and Tordo, 2001), which are now confirmed as reservoir hosts for 16 of the 18 recognized viruses (Kuzmin et al., 2011; Banyard et al., 2014; Rupprecht and Chikwamba, 2018). Rabies lyssavirus (RABV) is the most epidemiologically important and best studied virus in this group (Rupprecht and Chikwamba, 2018) and the only lyssavirus present in the Western Hemisphere (Rupprecht et al., 2011; Andres Velasco-Villa et al., 2017). Although reservoir hosts in North America are dominated by insectivorous bats and wild mesocarnivores (Messenger et al., 2002; Banyard et al., 2014), common vampire bats (*Desmodus rotundus*) are the primary reservoir for human and livestock rabies in the Neotropics (Schneider et al., 2009; Ulloa-Stanojlovic and Dias, 2020). Vampire bats feed primarily on mammal blood (Greenhall and Schmidt, 1988; Becker et al., 2018), facilitating RABV transmission to other species though saliva during biting (Aguilar-Setien et al., 2005; Johnson et al., 2014). Vampire bat RABV can be a significant public health burden via human exposures and fatalities, especially in resource-poor and communities (Condori-Condori et al., 2013; Stoner-Duncan et al., 2014; Fenton et al., 2020) and carries substantial economic costs via livestock mortality (Benavides et al., 2017).

Understanding how zoonotic viruses circulate and persist in their reservoir hosts is key to forecasting spillover risks and developing evidence-based interventions (Plowright et al., 2015; Plowright et al., 2019). RABV dynamics in vampire bats have been studied across parts of the species geographic range, providing important insights into viral transmission and maintenance. Work in Peru showed that vampire bat RABV does not follow density-dependent transmission, that bat culling activities predict greater viral exposure in bats, and that young bats are likely a critical demographic group for viral maintenance (Streicker et al., 2012). Longitudinal analyses of RABV antibody prevalence in Peru and French Guiana further suggested that viral persistence depends on vampire bat dispersal among sites and that most exposed bats likely gain transient immunity following exposure (i.e., without becoming infectious or experiencing mortality; Blackwood et al., 2013; de Thoisy et al., 2016). Spatiotemporal studies in Argentina and Peru showed that vampire bat RABV moves in traveling waves, likely driven by male-biased dispersal (Delpietro et al., 1972; Benavides et al., 2016; Streicker et al., 2016). Whether similar dynamics occur among vampire bats across Central America and the Caribbean remains a pressing and open question (Andres Velasco-Villa et al., 2017; Seetahal et al., 2018; Streicker et al., 2019).

Rabies is a notifiable disease in almost all countries of the Caribbean (CARICOM) (Seetahal et al., 2018), where it is listed as a priority zoonosis by the Caribbean Animal Health Network (Lefrancois et al., 2010). RABV is endemic in at least 10 CARICOM countries, of which vampire bats are the primary reservoir in French Guiana, Guyana, Suriname, Trinidad, and Belize (Seetahal et al., 2018). The latter is of particular interest given its proximity to other Central American countries such as Mexico and Guatemala, where vampire bat RABV transmission to livestock is common (Martínez-Burnes et al., 1997; Andrés Velasco-Villa et al., 2002, 2006; Ellison et al., 2014; Moran et al., 2015). Belize has reported rabies in domestic animals (i.e., cattle, sheep, and horses) across four of its six districts (World Organization for Animal Health, 2016), and outbreaks have increased in frequency, especially since 2016 (Fig. 1). RABV strains isolated from Belize livestock are the antigenic variants V-3 and V-11, both of which are vampire bat–associated variants first characterized in Mexico and Colombia (Andrés Velasco-Villa et al., 2002; Jiménez et al., 2017). Although human rabies has not been documented in Belize for over 20 years, vampire bat feeding on humans has been reported (McCarthy, 1989). Several vampire bats have tested positive for RABV in Belize (Constantine and Blehert, 2009; World Organization for Animal Health, 2016), underscoring the need for systematic data on spatial and temporal patterns of RABV in Belize vampire bat populations.

**Figure 1.**
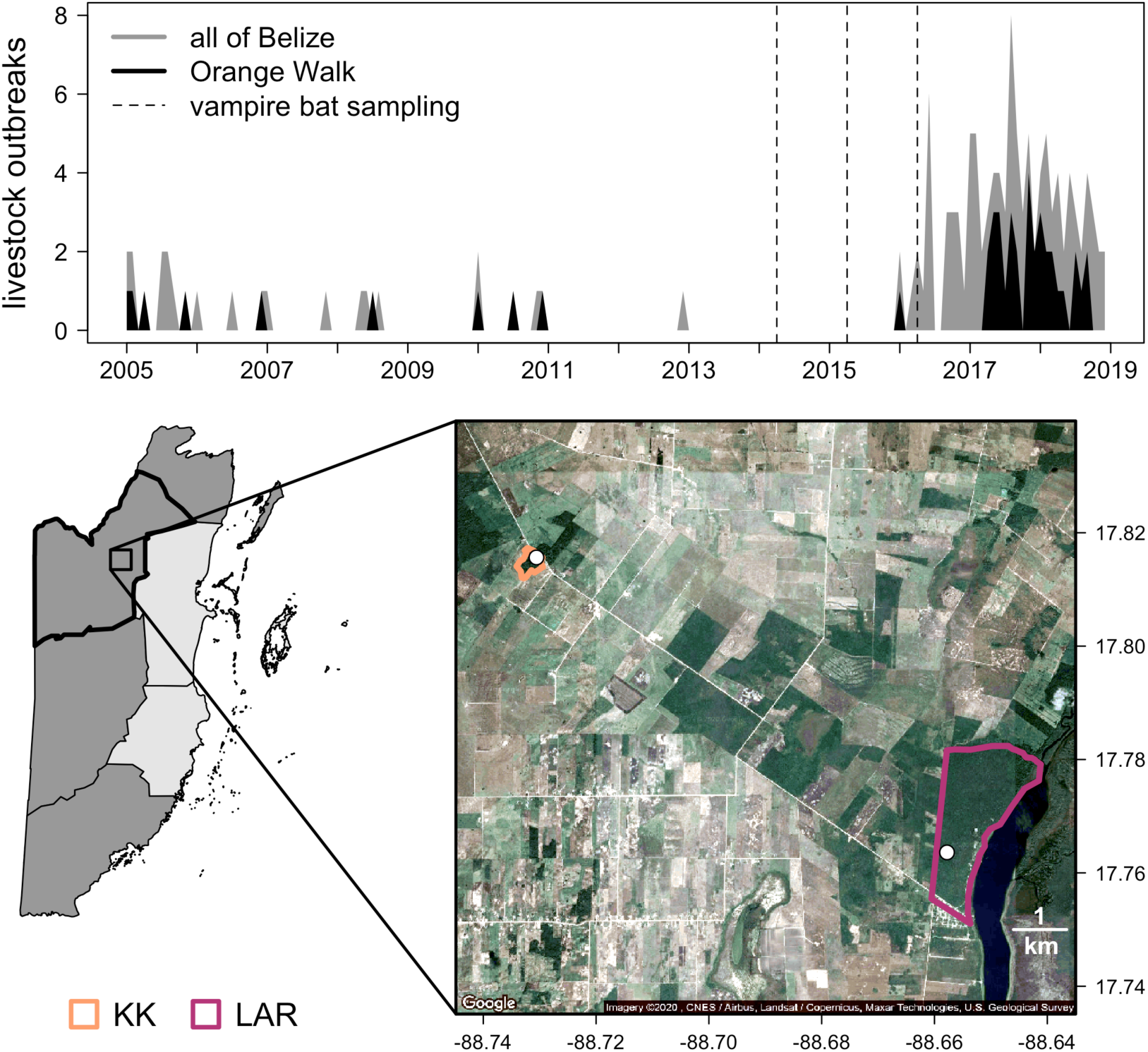
Temporal patterns of livestock rabies in Belize. (Upper) The number of outbreaks per month is shown in grey, with data from Orange Walk District in black. Data are derived from the World Organization for Animal Health. Dashed lines indicate vampire bat sampling events. (Lower) Map of Belize with districts with confirmed cases of livestock rabies displayed in grey and Orange Walk in black. The inset shows a satellite view of the study region: white and tan indicate agricultural and urban development, while dark green shows intact forest. Colored outlines show the two sites (KK and LAR), and sampled roosts are shown with white points.

To investigate patterns of vampire bat RABV over space and time in Belize, we performed a three-year serological study at two sites in Orange Walk District, where livestock rabies has increased in frequency (Fig. 1). In this agricultural region, cropland expansion in the 1990s, followed by heightened demand for livestock pasture in the 2000s, drove widespread deforestation (Patterson, 2016). These landscape changes fragmented previously intact forest and provide vampire bats with abundant livestock prey (Voigt and Kelm, 2006; Becker et al., 2018). We used serology to detect recent RABV exposure and population-level viral circulation in vampire bats, given that abortive RABV infections are common in bats and virus-neutralizing antibody (VNA) titers wane to undetectable levels within five months of exposure (Jackson et al., 2008; Turmelle et al., 2010). We first tested whether RABV circulates constantly or sporadically. Given past work on vampire bat rabies from Peru, we predicted RABV would display endemic persistence in Belize sites and that age would predict individual risk of RABV seropositivity (Streicker et al., 2012; Blackwood et al., 2013). Because we also found evidence for divergent RABV dynamics among sites, we used population genetics and mark–recapture data to ask if spatial patterns in virus exposure could be explained by inter-site bat dispersal.

## Materials and methods

### Vampire bat sampling

During 2014–2016, we sampled vampire bats in two areas of Orange Walk District: the Lamanai Archeological Reserve (LAR) and Ka’Kabish (KK). These sites are separated by 8 km of agricultural fields, secondary growth, and forest fragments and have been the basis of long-term bat research, representing contrasting community structure and land use (Fenton et al., 2001; Herrera et al., 2018). LAR contains 450 ha of semi-deciduous tropical forest at varying stages of regrowth after clearing for settlement and agriculture from the Maya period onward. Habitats range from closed canopy forest to clearings, and the protected reserve is bordered by the New River Lagoon, forest, and agricultural fields. KK is a smaller fragment (approximately 45 ha) of secondary forest overlaying another Maya city, surrounded by agriculture and pasture (Fig. 1).

We sampled bats during 1–2 weeks annually between late April and early May (Table S1), which coincides with the transition between the dry and wet seasons in Belize. Daily high temperatures at both sites were 29–37 °C with sunny days and occasional afternoon or evening thunderstorms. We targeted single known roosts per site and used mist nets (6, 9, and 12 meters) at the entrances of both roosts (covering the only exit for the LAR roost and both main exits for the KK roost) as well as along known flight paths from approximately 19:00 to 22:00. We also set an Austbat harp trap (Titley Scientific, capture area of approximately 2 meters by 1.8 meters) from 18:00 until 5:00 on different nights per site to maximize capture. Mist nets were monitored continuously, and harp traps were checked every half hour until midnight and again before dawn. Because most captures were obtained via mist nets, bats were primarily captured leaving roosts. However, we note that other roosts do exist in the broader study region, and thus our capture efforts only sampled a subset of the broader vampire bat metapopulation (Trajano, 1996).

All vampire bats (*n*=122) were placed in individual holding bags upon capture and issued a uniquely numbered incoloy wing band (3.5 mm, Porzana Inc.). We classified age as sub-adult or adult based on fusion of phalangeal epiphyses (Delpietro and Russo, 2002). Reproductive status was determined by the presence of scrotal testes in males and pregnancy or lactation in females. Across all three years, both sites had uneven sex composition, with KK having a more skewed sex ratio (86% males) than LAR (65% males). From a subset of bats, we obtained blood samples by lancing the propatagial vein with 23-gauge needles, followed by collection with heparinized capillary tubes. We obtained serum (*n*=67) by centrifuging blood in serum separator tubes. To assess population genetic connectivity, we collected two 2 mm wing biopsy punches stored in 95% ethanol. Samples were frozen at –20°C until long-term –80°C storage. Field protocols followed guidelines for safe and humane handling of bats issued by the American Society of Mammalogists (Sikes et al., 2016) and were approved by the University of Georgia Animal Care and Use Committee (A2014 04-016-Y3-A5). Bat sampling was authorized by the Belize Forest Department under permits CD/60/3/14(27), CD/60/3/15(21), and WL/1/1/16(17).

### Rabies serology

Serum was heat inactivated at 56°C for one hour before exportation to the MRC–University of Glasgow Centre for Virus Research. To detect VNA, we used a modified pseudotype virus micro-neutralization test (Meza et al., 2020). Specifically, we used a pseudotype virus with a murine leukemia virus backbone and a RABV glycoprotein gene (strain CVS-11) rather than pathogenic RABV (Kuzmin et al., 2008; Streicker et al., 2012). Along with each batch of samples, we tested a dilution series with known concentrations of standard rabies immunoglobulin (SRIG) from 0.01 to 0.2 International Units per mL (IU/mL). In each test well, which contained a dilution of serum and challenge virus, the number of fluorescing cells (i.e., virus-infected cells represented by an accumulation of pixels in a binary image) was calculated across five image fields using ImageJ (Abràmoff et al., 2004). We next fit a binomial (logit link) generalized linear mixed model (GLMM) to the SRIG data, considering titers > 0.1 IU/mL as positive and titers < 0.1 IU/mL as negative. The fixed effect was the count of virus-infected mouse neuroblastoma (N2A) cells under the 1:10 dilution, and we included a random intercept of assay date. After fitting this GLMM to the SRIG data, we predicted the probability that field-collected samples were seropositive, using the fluorescing cell counts of the 1:10 serum dilution (Fig. S1). We considered bats with mean predicted probabilities (averaged over the five microscopy fields) greater than 95% to be seropositive. We then used a secondary GLMM with a log-normal distribution (fit to the full concentration series of SRIG 0.1–0.2 IU/mL, 1:10 dilution) to predict VNA titers, which we used to evaluate whether seropositive individuals had predicted titers far above our VNA cutoff, which would indicate a low risk of false positives (Fig. S2). GLMMs were fit using the *lme4* package in R (Bates et al., 2015).

### Statistical analysis of serological data

We also used the *prevalence* package in R to estimate RABV seroprevalence and 95% confidence intervals (*i*) across all data and (*ii*) per year per site. To quantify the degree of synchrony in RABV seroprevalence between sites, we estimated the zero-lag cross-correlation coefficient (*r*; –1 to 1) between the three-year serological time series (Bjørnstad et al., 1999).

To test the spatial, temporal, and individual predictors of RABV seropositivity, we fit a set of GLMs with binomial errors and a logit link. GLMs included year, site, sex, reproductive status, age, and the interaction between year and site; we did not fit a single GLM with all predictors to prevent overfitting. Owing to a low number of recaptured bats with sera tested for RABV VNA on multiple occasions (*n*=3), we randomly selected one capture event for these individuals in our GLMs rather than include bat identification number as a random effect. We used the *MuMIn* package and Akaike information criterion corrected for small sample size (AICc) to compare candidate GLMs fit to a dataset free of missing values (*n*=61), and models within two ΔAICc were considered competitive (Burnham and Anderson, 2002; Barton, 2013). We assessed fit (*R*^*2*^) using the *rsq* package (Nagelkerke, 1991). Because male behavior and disperser bias could affect rabies exposure (Streicker et al., 2016; Huguin et al., 2018), we also tested for a correlation between the site- and year-specific fractions of males and seroprevalence.

### Population connectivity

Vampire bats often use multiple roosts (Wilkinson, 1985), and inter-roost movement typically occurs within a relatively small area (i.e., from 2–10 km) for both sexes, although some male bias is evident (Trajano, 1996; Delpietro et al., 2017). However, dispersal over longer distances is almost exclusively undertaken by males (Greenhall et al., 1983; Streicker et al., 2016). Bat contact at shared feeding grounds within the foraging range of our sites could also facilitate inter-site virus transmission (Greenhall et al., 1971; Blackwood et al., 2013). To first assess connectivity through bat population genetics, we extracted DNA from wing biopsies (LAR=41, KK=28) using DNeasy Tissue Kits (Qiagen) and amplified nine previously developed nuclear microsatellite loci using a Multiplex PCR Kit (Qiagen; Piaggio et al., 2008). Microsatellites are biparentally inherited markers that have previously captured patterns of vampire bat population structure at both large and fine spatial scales and correlate with patterns of rabies virus spread (Streicker et al., 2016; Huguin et al., 2018). We used Geneious (BioMatters) for microsatellite scoring (Kearse et al., 2012). In R, we next used the *poppr* and *hierfstat* packages to process microsatellite data and assess population structure (Goudet, 2005; Kamvar et al., 2014). We estimated genetic distance as Nei’s pairwise *F*_*ST*_ (Goudet et al., 1996) and visualized population structure using a principal components analysis (PCA).

As a secondary method, we applied mark–recapture analysis to our field sampling data (2014–2016) and to later vampire bat sampling efforts (2017–2019) to estimate the relative frequency of inter-site movement. We used the *prevalence* package to estimate dispersal events relative to all recaptures and the associated 95% confidence interval. To test if males are less roost faithful than females, we used a GLM with sex as the predictor and inter-roost movement as a binomial response. We also used a GLMM with a random effect of identification number to assess sex biases in annual recapture (i.e., binomial response) of all banded individuals, as lack of recapture suggests mortality or dispersal. We did not use mark–recapture to estimate colony size owing to limited recaptures in 2014–2016, when serology data were available, and to weak evidence for density-dependent RABV transmission in vampire bats (Streicker et al., 2012).

## Results

We found that 47.8% of the 67 vampire bats assayed for RABV VNA in northern Belize were seropositive across sites and years (95% CI: 36.25–59.51%). Overall seroprevalence was similar between KK and LAR (44.1% and 51.5%; Fig. 2A), with most inter-site variation in 2015 (KK=29.4%, LAR=80%). Estimated VNA titers for seropositive bats were generally well above the pre-defined threshold for positivity across all sites and years, with the exception of LAR in 2014 (Fig. S2). Correlation between annual seroprevalence in KK and LAR was negative at the zero lag (*r*_0_=–0.68) and suggested a phase lag of at least a year (Fig. 2A). Model comparison did not identify site, year, or individual predictors of seropositivity (Table 1). For the three recaptured bats tested for VNA (all male), we detected one seroconversion from negative to positive and another from positive to negative, both associated with large titer changes (Fig. S3). The proportion of males per site and year did not correlate with seroprevalence (*r*=0.29, *p*=0.57).

**Table 1.** Candidate GLMs predicting vampire bat RABV seropositivity in Belize. Competing models are ranked by ΔAICc with the number of coefficients (*k*) Akaike weights (*w*_*i*_), and *R*^*2*^.

| GLM structure | $k$ | $\Delta\text{AICc}$ | $w_i$ | $R^2$ |
| --- | --- | --- | --- | --- |
| Seropositivity ~ 1 | 1 | 0 | 0.21 | 0 |
| Seropositivity ~ site | 2 | 1.33 | 0.11 | 0.02 |
| Seropositivity ~ sex | 2 | 1.46 | 0.1 | 0.01 |
| Seropositivity ~ age | 2 | 1.93 | 0.08 | 0 |
| Seropositivity ~ reproduction | 2 | 2.13 | 0.07 | 0 |
| Seropositivity ~ reproduction + sex | 3 | 2.71 | 0.05 | 0.04 |
| Seropositivity ~ sex + site | 3 | 3.25 | 0.04 | 0.02 |
| Seropositivity ~ age + site | 3 | 3.29 | 0.04 | 0.02 |
| Seropositivity ~ reproduction + site | 3 | 3.45 | 0.04 | 0.02 |
| Seropositivity ~ age + sex | 3 | 3.53 | 0.04 | 0.02 |
| Seropositivity ~ year * site | 6 | 3.74 | 0.03 | 0.16 |
| Seropositivity ~ year | 3 | 4.11 | 0.03 | 0.01 |
| Seropositivity ~ age + reproduction | 3 | 4.15 | 0.03 | 0 |
| Seropositivity ~ reproduction + sex + site | 4 | 4.62 | 0.02 | 0.04 |
| Seropositivity ~ age + reproduction + sex | 4 | 5 | 0.02 | 0.04 |
| Seropositivity ~ age + sex + site | 4 | 5.34 | 0.01 | 0.03 |
| Seropositivity ~ age + reproduction + site | 4 | 5.54 | 0.01 | 0.02 |
| Seropositivity ~ year + sex | 4 | 5.74 | 0.01 | 0.02 |
| Seropositivity ~ year + site | 4 | 5.74 | 0.01 | 0.02 |
| Seropositivity ~ age + year | 4 | 6.22 | 0.01 | 0.01 |
| Seropositivity ~ year + reproduction | 4 | 6.38 | 0.01 | 0.01 |
| Seropositivity ~ year + reproduction + sex | 5 | 7.1 | 0.01 | 0.04 |
| Seropositivity ~ year + sex + site | 5 | 7.79 | 0 | 0.03 |
| Seropositivity ~ age + year + site | 5 | 7.86 | 0 | 0.03 |
| Seropositivity ~ age + year + sex | 5 | 7.97 | 0 | 0.02 |
| Seropositivity ~ year + reproduction + site | 5 | 8.04 | 0 | 0.02 |
| Seropositivity ~ age + year + reproduction | 5 | 8.6 | 0 | 0.01 |

**Figure 2.**
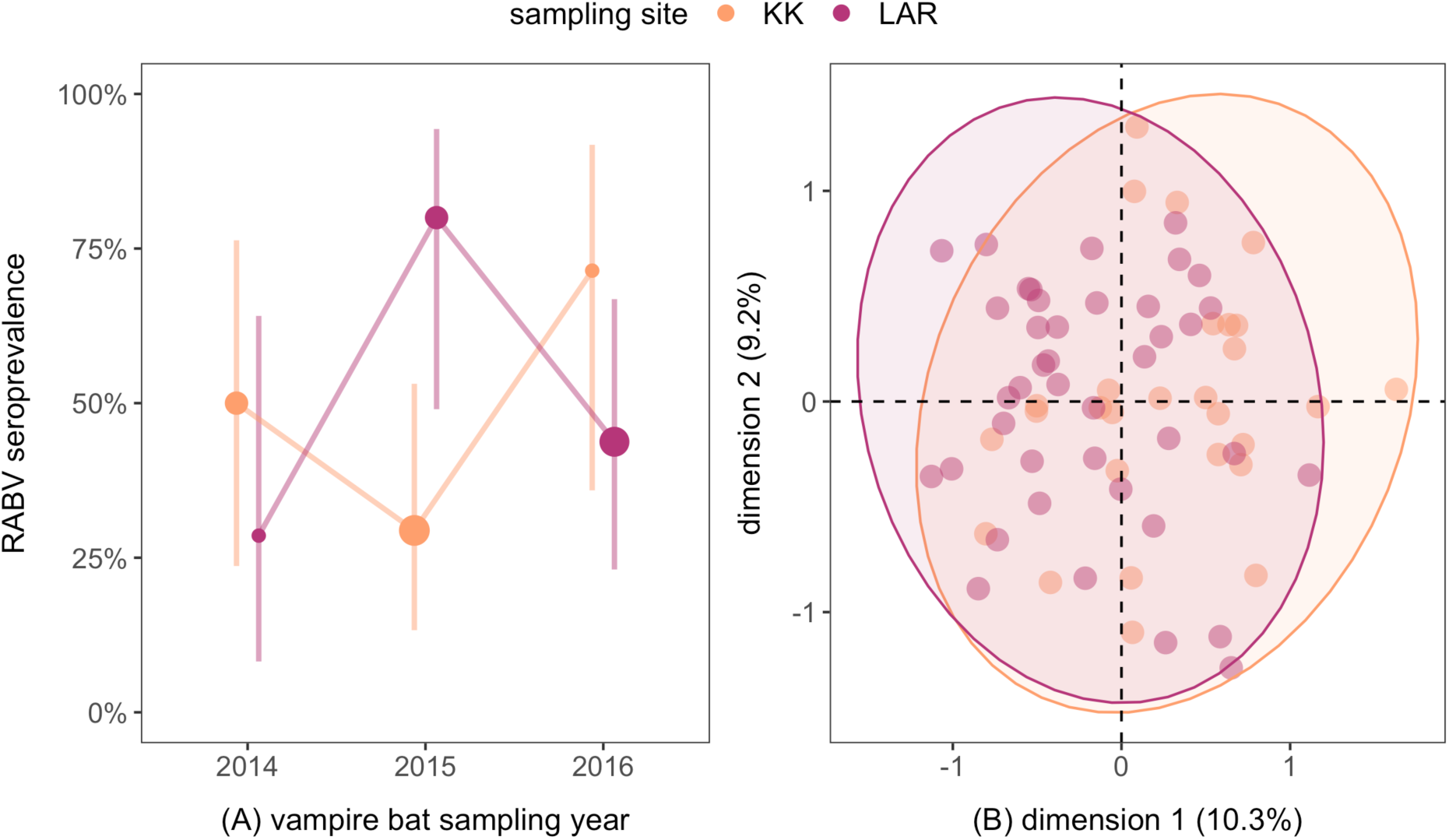
(A) RABV seroprevalence across the three study years, colored by site (KK and LAR) and scaled by sample size. Vertical lines show the 95% confidence intervals (Wilson interval). (B) PCA of microsatellite data for nine loci (*n*=69 individuals, 64 alleles). Shown are the first two axes of the PCA with points colored by site and grouped using concentration ellipses.

Analyses of vampire bat population structure suggested very low genetic differentiation between sites (*F*_*ST*_=0.015), and the PCA also suggested general panmixia (Fig. 2B). Consistent with population genetic results, mark–recapture data across a broader sample of 75 recaptured bats (*n*=193 captures into 2019) revealed rare but detectable inter-year movements between sites, including two males and one female (4/118 recaptures, 95% CI: 1.3–8.4%; Fig. 3). Perhaps unsurprisingly given these relatively rare inter-site dispersals, males were not significantly more likely to move between sites (OR=2.24, *p*=0.51), although one adult male was recaptured between sites twice from 2016 to 2019 (LAR to KK to LAR). Lastly, when considering the likelihood of recapturing a banded bat annually (*n*=234 individuals over five years), males had weakly lower odds of annual recapture (11%) compared to females (23%; OR=0.23, *p*=0.1)

**Figure 3.**
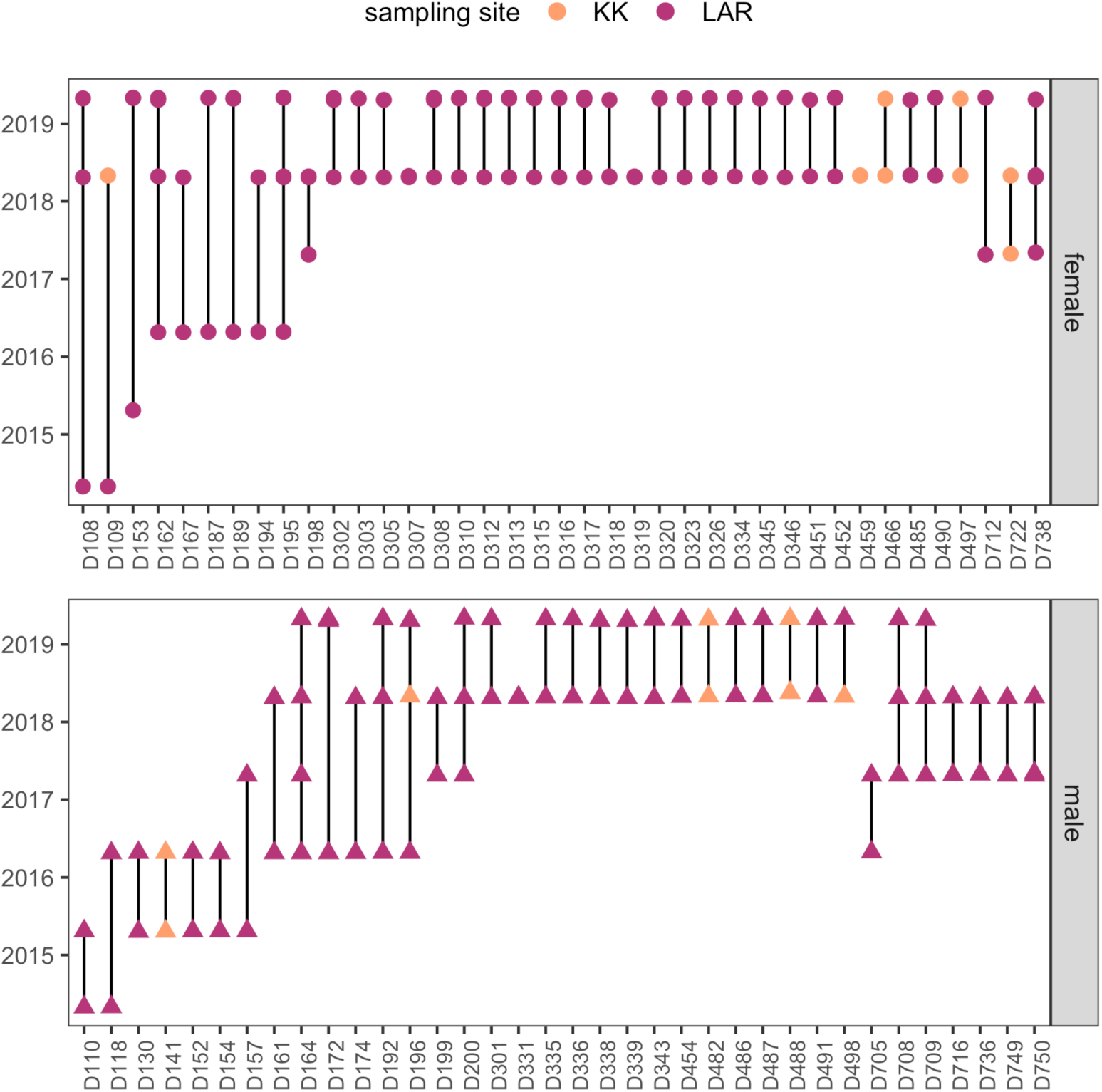
Roost fidelity of recaptured vampire bats banded during 2014–2018 and sampled during 2015–2019, colored by site and stratified by sex. Points with no lines are individuals recaptured in the same year; no individuals were recaptured in different sites in the same year.

## Discussion

This study provides results of the first active surveillance for RABV circulation among vampire bats in Belize. We show RABV is actively circulating in vampire bats in the north of Belize and that the virus appears to show endemic dynamics. Between two sites, seroprevalence was high, inversely related, and phase lagged by likely a year. Microsatellite data demonstrated substantial gene flow between sites, and mark–recapture analyses detected rare inter-site bat movements. Together, these observations suggest one plausible explanation for the inverse seroprevalence dynamics could be infrequent but ecologically relevant vampire bat movement among sites.

The detection of rabies VNA demonstrates active viral circulation in vampire bats in Belize, consistent with RABV circulation in *Desmodus rotundus* in adjacent Guatemala and Mexico (Andrés Velasco-Villa et al., 2002, 2006; Ellison et al., 2014). However, our average estimate of RABV seroprevalence among bats in Belize (47.8%) was notably higher than for vampire bats sampled in other regions, including Peru (10.5%; Streicker et al., 2012), Brazil (7.3% and 12.2%; Langoni et al., 2008; Almeida et al., 2011), French Guiana (14.2%; de Thoisy et al., 2016), and Guatemala (9%; Ellison et al., 2014). We used a higher (more stringent) VNA cutoff for seropositivity (0.1 IU/mL) than some other studies of bat rabies (e.g., 0.05 and 0.06 IU/mL; Streicker et al., 2013; Ellison et al., 2014), and lower VNA cutoffs would only increase our seroprevalence estimate. Given ongoing rabies outbreaks in Belize, sampling bats during the epizootic could have facilitated a higher likelihood of detecting seropositive bats, as VNAs from recent viral exposures would not yet have waned to undetectable levels (Jackson et al., 2008; Plowright et al., 2019). Vampire bats in our sites also co-roost with other bat species, including *Glossophaga soricina* and *Trachops cirrhosus* (Fenton et al., 2001), which have tested positive for RABV VNA in other countries, so exposures to non-vampire bat RABV variants could have elevated seroprevalence (Ellison et al., 2014; Almeida et al., 2019). More broadly, the high frequency of seropositive individuals here provides further evidence that many apparently healthy bats are exposed to RABV and can clear peripheral viral infection without showing disease (Turmelle et al., 2010; Streicker et al., 2012; Ellison et al., 2014; de Thoisy et al., 2016).

Host populations connected by movement should spatially homogenize pathogen transmission (Hess, 1996). Prevalence of RABV exposure in vampire bats did not differ significantly between our two sites. However, site-specific seroprevalence estimates were negatively associated at the zero-year lag, implying localized RABV dynamics are out of phase. More years of serological data are needed to confirm such patterns, but one explanation for this inverse relationship could be bat movement that causes traveling waves of virus transmission (Bjørnstad et al., 1999). Our microsatellite data suggest historic panmixia of bats in this region of Belize, consistent with low population structure at small spatial scales in Mexico, Peru, and French Guiana, as also inferred using nuclear markers (Romero-Nava et al., 2014; Streicker et al., 2016; Huguin et al., 2018). Additionally, mark–recapture detected rare but contemporary annual movement between sites, an 8 km distance relative to mean home ranges of 2–3 km (Burns and Crespo, 1975; Trajano, 1996). The relatively low frequency of inter-site movement could stem from recent habitat fragmentation (Ingala et al., 2019) or the presence of additional unsampled roosts in the broader landscape (Trajano, 1996); in 2019, the authors photographed banded vampire bats in a newly detected cave roost approximately 6 km from the KK and LAR roosts. Additionally, inter-site movement over larger spatial scales has been shown to be more frequent in males (Delpietro et al., 2017), and prior population genetics analyses suggest skewed sex ratios could facilitate spatial spread of RABV by male-biased dispersal (Streicker et al., 2016). Although we did not detect a sex difference in the frequency of inter-site movement, male bats were somewhat less likely to be recaptured between years, which could indicate movement to other roosts in the landscape. The stronger male bias in KK could also explain the phase lag in seroprevalence if this site sources more infected dispersers (Becker, Snedden et al., 2018).

Fine-scale studies of bat movement are needed to better quantify contemporary connectivity and to understand how alternative roosts and male dispersal affect RABV dynamics. Additionally, use of mitochondrial markers could also help illuminate how sex-dependent behavior shapes virus transmission (Streicker et al., 2016; Huguin et al., 2018). More broadly, however, our results suggest that the degree of connectivity found here likely enables virus persistence through inter-site contacts but is not sufficient to synchronize infection dynamics across sites (Blackwood et al., 2013). In further support of this idea, the inverse relationship in seroprevalence between our Belize sites (<10 km) is consistent with prior rates of RABV dispersal in vampire bats of 9–17 km per year (Benavides et al., 2016; Streicker et al., 2019).

Future phylogenetic analyses of RABV isolates from across Belize as well as nearby countries could help differentiate hypotheses of a persistent transmission cycle in Belize against the potential role of viral invasions and spatial spread (Andrés Velasco-Villa et al., 2006; Biek et al., 2007; Streicker et al., 2019). Although our serological data are consistent with endemic maintenance of RABV in local vampire bat populations, livestock rabies outbreaks are known to show epidemic dynamics, including in Belize, where outbreaks were not reported in livestock in 2014 and 2015 (World Organization for Animal Health, 2016). Vaccination of livestock could limit outbreak detection or cause apparent viral extinctions. Future diet analyses of vampire bats could also address whether the mismatch between observed seroprevalence in vampire bats and outbreaks in livestock reflects temporal changes in contact rates between the reservoir hosts and recipient hosts (e.g., large temporal shifts toward feeding on livestock prey). The latter could be especially informative given intensifying agricultural activity in this region (Patterson, 2016), which could affect both bat feeding patterns and local movement (Botto Nuñez et al., 2020).

Despite active circulation of RABV in Belize vampire bats and associated rabies outbreaks in livestock, we caution that culling of vampire bats through destruction of roosts or application of anticoagulant paste is unlikely to eliminate rabies (Linhart et al., 1972; Streicker et al., 2012; Johnson et al., 2014). Moreover, theory suggests that culling can increase RABV transmission by increasing dispersal of infected bats across larger spatial scales (Blackwood et al., 2013). Destruction of roosts also affects population viability of other bat species that reside with vampire bats, threatening ecosystem services provided by insectivorous and frugivorous bats (O’Shea et al., 2016). Although we are unaware of culling applied to roosts in our sites, vampire bat control has been applied elsewhere in Belize, and landowners maintain negative perceptions of bats (Shapiro et al., 2020). Alternative interventions, such as vaccination of humans and livestock, efforts to reduce bite incidence, and oral vaccination of bats themselves, could be more sustainable and would reduce negative impacts of rabies control on Neotropical bat communities (Anderson et al., 2014; Stoner-Duncan et al., 2014; Bakker et al., 2019).

## Data accessibility

Individual-level RABV serological data are available in Mendeley Data (Becker et al., 2020).

## Acknowledgements

For coordinating vampire bat sampling and permits, we thank Neil Duncan, Mark Howells, Venetia Briggs, and staff of Lamanai Field Research Center. We also thank the many colleagues who helped capture bats during our annual surveys in Belize. We thank two anonymous reviewers for constructive feedback on this manuscript.

