## Supplemental Material for "Temporal patterns of vampire bat rabies and host connectivity in Belize"

### Supplemental Information

Daniel J. Becker, Alice Broos, Laura M. Bergner, Diana K. Meza, Nancy B. Simmons, M. Brock Fenton, Sonia Altizer, Daniel G. Streicker

**S1. Capture details**

**S2. Rabies seropositivity**

**S3. Recaptured bats**

**S1. Capture details**

Table S1. Capture numbers (stratified by sex) per site and date during the serological sampling years (2014-2016). Sex information is missing for a small number of these captures (*n*=4).

| Site | Date | Total | Females | Males |
|---|---|---|---|---|
| KK | 4/29/2014 | 6 | 0 | 6 |
| | 4/30/2014 | 5 | 1 | 4 |
| | 5/1/2014 | 8 | 1 | 7 |
| | 5/2/2014 | 2 | 1 | 0 |
| | 5/3/2014 | 2 | 1 | 1 |
| | 4/22/2015 | 14 | 2 | 12 |
| | 4/23/2015 | 3 | 0 | 3 |
| | 4/27/2016 | 10 | 1 | 9 |
| LAR | 4/30/2014 | 4 | 1 | 3 |
| | 5/1/2014 | 4 | 2 | 2 |
| | 5/2/2014 | 1 | 0 | 1 |
| | 5/3/2014 | 3 | 1 | 2 |
| | 4/21/2015 | 2 | 0 | 2 |
| | 4/23/2015 | 3 | 1 | 2 |
| | 4/24/2015 | 8 | 2 | 6 |
| | 4/25/2016 | 17 | 5 | 12 |
| | 4/26/2016 | 6 | 2 | 4 |
| | 4/27/2016 | 15 | 4 | 9 |
| | 4/29/2016 | 9 | 6 | 2 |

**S2. Rabies seropositivity**

Figure S1. Raw counts of virus-infected cells from the RFFIT assay in relation to individual bat predicted probability of being seropositive (from the SRIG binomial GLMM). Individuals with predicted probabilities greater than 95% were considered seropositive for RABV VNA.

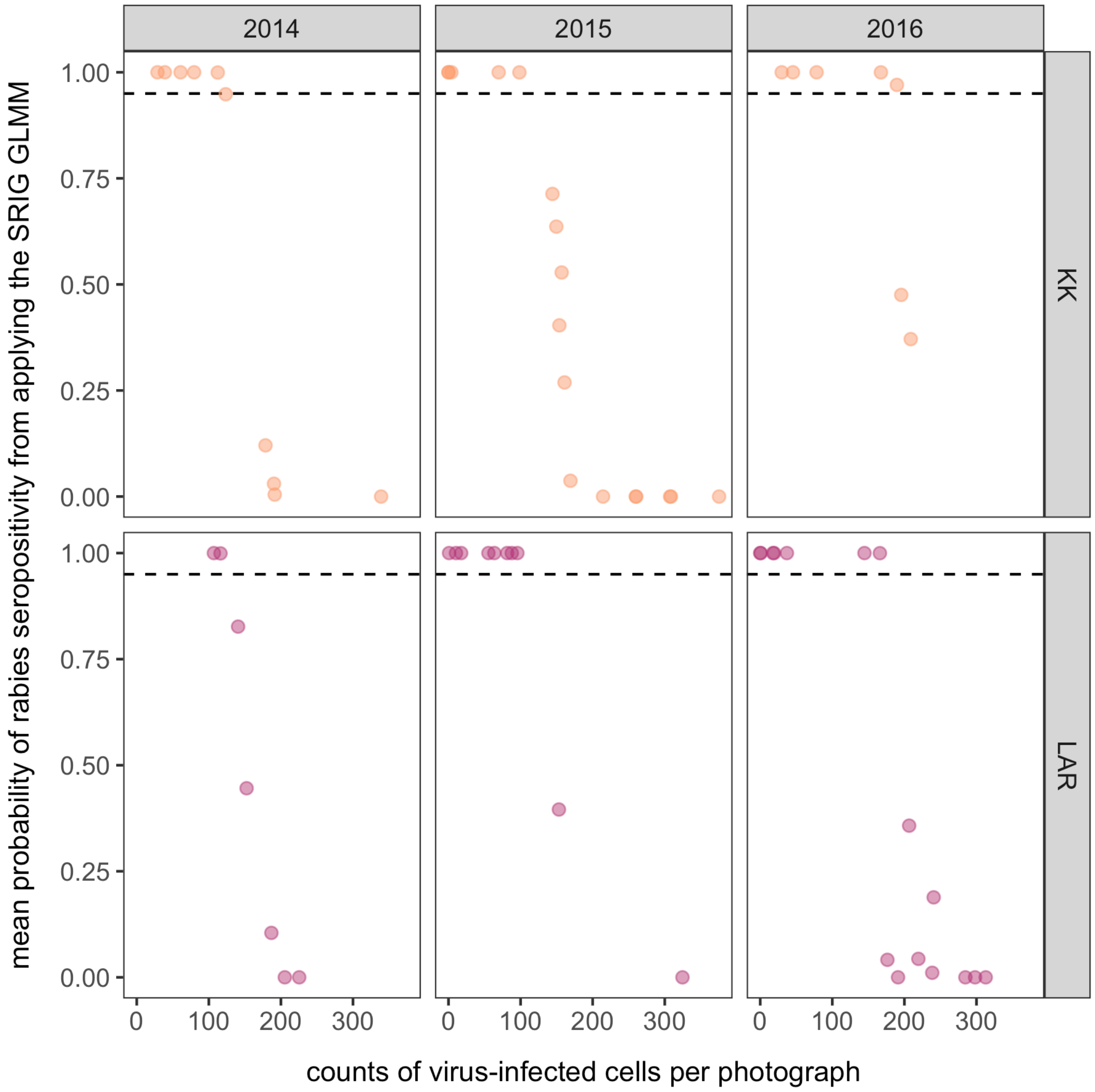

Figure S2. Relationship between predicted VNA titers (from the SRIG log-normal GLMM) and predicted serological status (from the SRIG binomial GLMM, jittered to reduce overlap) for each site and year. The dashed line shows the serological cutoff used in the binomial SRIG GLMM.

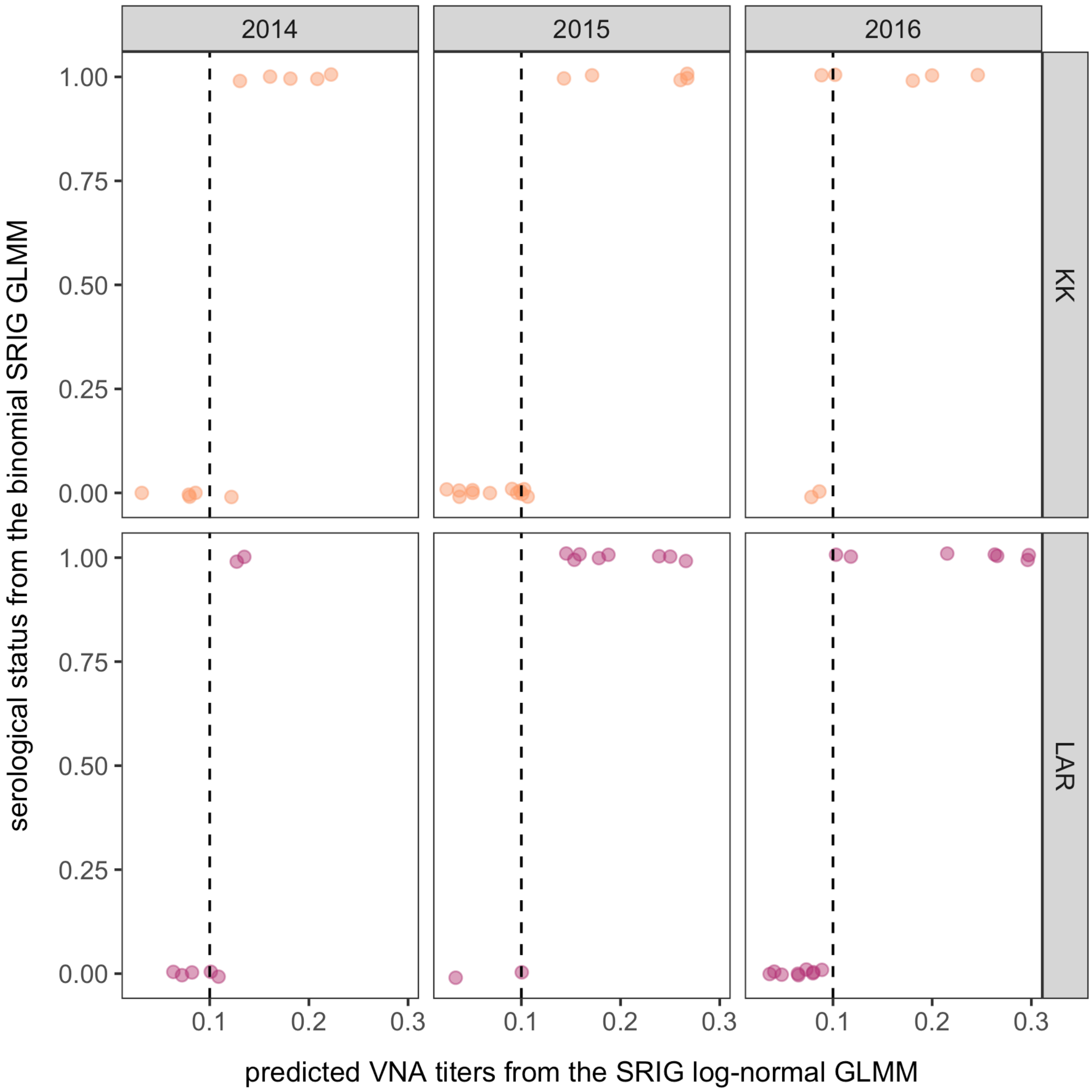

### S3. Recaptured bats

Figure S3. Changes in RABV serological status and predicted VNA titers between years for recaptured bats.

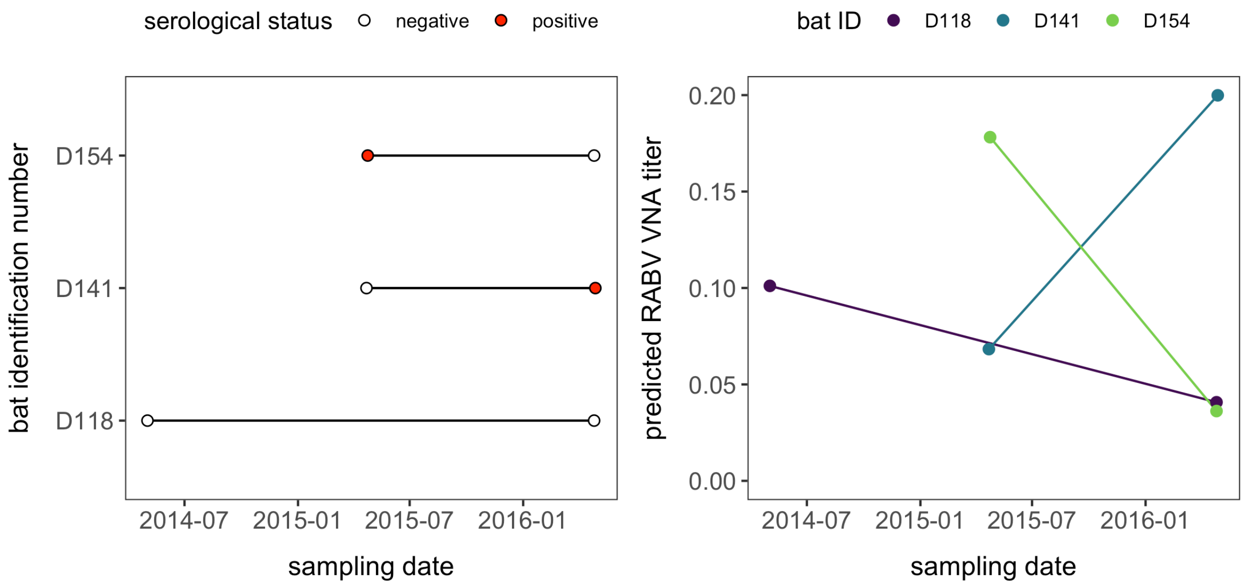
